# Evaluating spatial and network properties of NMDA-dependent neuronal connectivity in mixed cortical cultures

**DOI:** 10.1101/2022.03.10.483856

**Authors:** Catherine P. Rojvirat, Joshua R. Berlin, Tuan D. Nguyen

## Abstract

A technique combining fluorescence imaging with Ca^2+^ indicators and single-cell laser scanning photostimulation of caged glutamate (LSPS) allows identification of functional connections between individual neurons in mixed cultures of rat neocortical cells as well as observation of synchronous spontaneous activity among neurons. LSPS performed on large numbers of neurons yielded maps of functional connections between neurons and allowed calculation of neuronal network parameters. LSPS also provided an indirect measure of excitability of neurons targeted for photostimulation. By repeating LSPS sessions with the same neurons, stability of connections and change in the number and strength of connections were also determined. Experiments were conducted in the presence of bicuculline to study the properties of excitatory neurotransmission. The AMPA receptor inhibitor, 6-Cyano-7-nitroquinoxaline-2,3-dione (CNQX), abolished synchronous neuronal activity but had no effect on connections mapped by LSPS. In contrast, the NMDA receptor inhibitor, 2-Amino-5-phosphono-pentanoic acid (APV), dramatically decreased the number of functional connections between neurons while also affecting synchronous spontaneous activity. Functional connections were also decreased by increasing extracellular Mg^2+^ concentration. These data demonstrated that LSPS mapping interrogates NMDA receptor-dependent connectivity between neurons in the network. A GluN2A-specific inhibitor, NVP-AAM077, decreased the number and strength of connections between neurons as well as neuron excitability. Conversely, the GluN2A-specific positive modulator, GNE-0723, increased these same properties. These data showed that LSPS can be used to directly study perturbations in the properties of NMDA receptor-dependent connectivity in neuronal networks. This approach should be applicable in a wide variety of *in vitro* and *in vivo* experimental preparations.

## 1. Introduction

First proposed by Donald Hebb, the “cell-assembly” hypothesis (Hebb, 1949), in which groups of neurons firing together can form the basis of functional states in the brain, has generated renewed interest (Harris, 2005; Huyck & Passmore, 2013; Plenz & Thiagarajan, 2007; Wallace, 2010). Indeed, dynamic changes at the network level may help define sleep/wake states, perception, cognition, and memory formation (Carrillo-Reid & Yuste, 2020a). Moreover, this hypothesis, if true, raises the question of what precisely constitutes the fundamental building block of neuronal computation (Carrillo-Reid & Yuste, 2020b).

Answering this question experimentally by probing and recording neuronal activity at the network level remains a daunting challenge (Harris, 2005; Yuste, 2015). While fMRI and PET scans are useful for defining the macroscopic functional architecture of brain networks (Shumskaya et al., 2012), they lack the spatial and temporal resolution to explore the microcircuitry. Whole-cell patch clamping using multiple electrodes has sufficient resolution but is usually limited to just a few neurons (Perin and Markram, 2013). When combined with laser scanning photostimulation (Katz & Dalva, 1994), whole-cell patch clamping was highly successful in understanding microcircuits in brain slice preparations (Shepherd & Svoboda, 2005; Sturm et al., 2014), even though recording of activity was still limited to a single neuron.

Recently, laser scanning photostimulation (LSPS) has been combined with Ca^2+^ imaging (Nguyen et al., 2017) to allow for stimulation of and simultaneous recording of activity from a large population of neurons (~100). This technique can be used to understand the functional connectivity of the network at the single-cell level. Moreover, recent advances in molecular and genetic tools (Packer et al., 2013; Wang et al., 2019) can be incorporated into this technique to look at neuronal networks that contain multiple cell types and/or complex macro- and microcircuitry architectures.

This report examines how the combination of LSPS and Ca^2+^ imaging can be used to study functional changes in neuronal networks brought about by altering individual functional connections between neurons within the network. While the present study was carried out with mixed glial/neuronal cultures, the approach outlined here could be utilized for *in vitro* and *in vivo* studies and, therefore, have broad applicability to understand the cellular basis for changes in network function in the brain.

## 2. RESULTS

### 2.1. Identifying location and activity of individual neuronal soma during spontaneous synchronous activity in mixed neuronal cultures

Mixed cortical cultures contain neurons as well as glia. In our mixed cultures, 46.3±1.8% (mean ± S.E.M., n=7) of the cells counted by DAPI staining were also identified as neurons by immunohistological staining with a MAP2 antibody (not shown). The remaining cells were presumably glia, consistent with additional staining using antibodies against GFAP (for astrocytes) and IBA1 (for microglia).

Periodic elevations of intracellular Ca^2+^ have often been used as an indirect indicator of neuronal activity. However, both neurons and glia produce elevations in intracellular Ca^2+^ as part of their cellular signaling. For this reason, an approach to distinguish neuronal from non-neuronal Ca^2+^ signaling was necessary. To achieve this goal, we took advantage of the morphological differences between the cell bodies of glia and the soma of neurons as well as the observation that cortical neurons in culture form functional networks. These networks are characterized by spontaneous neuronal electrical activity that is periodic and synchronized, and likewise produces synchronous increases in intracellular Ca^2+^ among interconnected neurons. Glial cells, such as astrocytes, did not participate in these types of synchronous events but tended to have longer periods of elevated Ca^2+^, commonly referred to as calcium waves (Guthrie et al., 1999). These factors were used to identify the location of soma and activity of individual neurons with the procedure described below.

A brightfield image was first processed to detect and record XY-coordinates of cell body centroids in order to define regions of interest (ROIs) around the centroids (Figure 1A). These steps were part of an automated ROI detection macro written in ImageJ. At the working magnification of our system with a 4x objective, only neuronal somas were routinely detected. Calculation of ∆F/F for each ROI was performed within a 7×7 pixel area (approximately 20 µm across) centered on each cell, where baseline fluorescence intensity (F) was the average fluorescence intensity of the ROI over the observation period. As a visual check during validation of this program, calculated ROIs were superimposed on brightfield images (Figure 1B) and images recorded at peak fluorescence intensity (Figure 1C) to ensure visual center-alignment of ROIs with soma positions.

**Figure 1.**
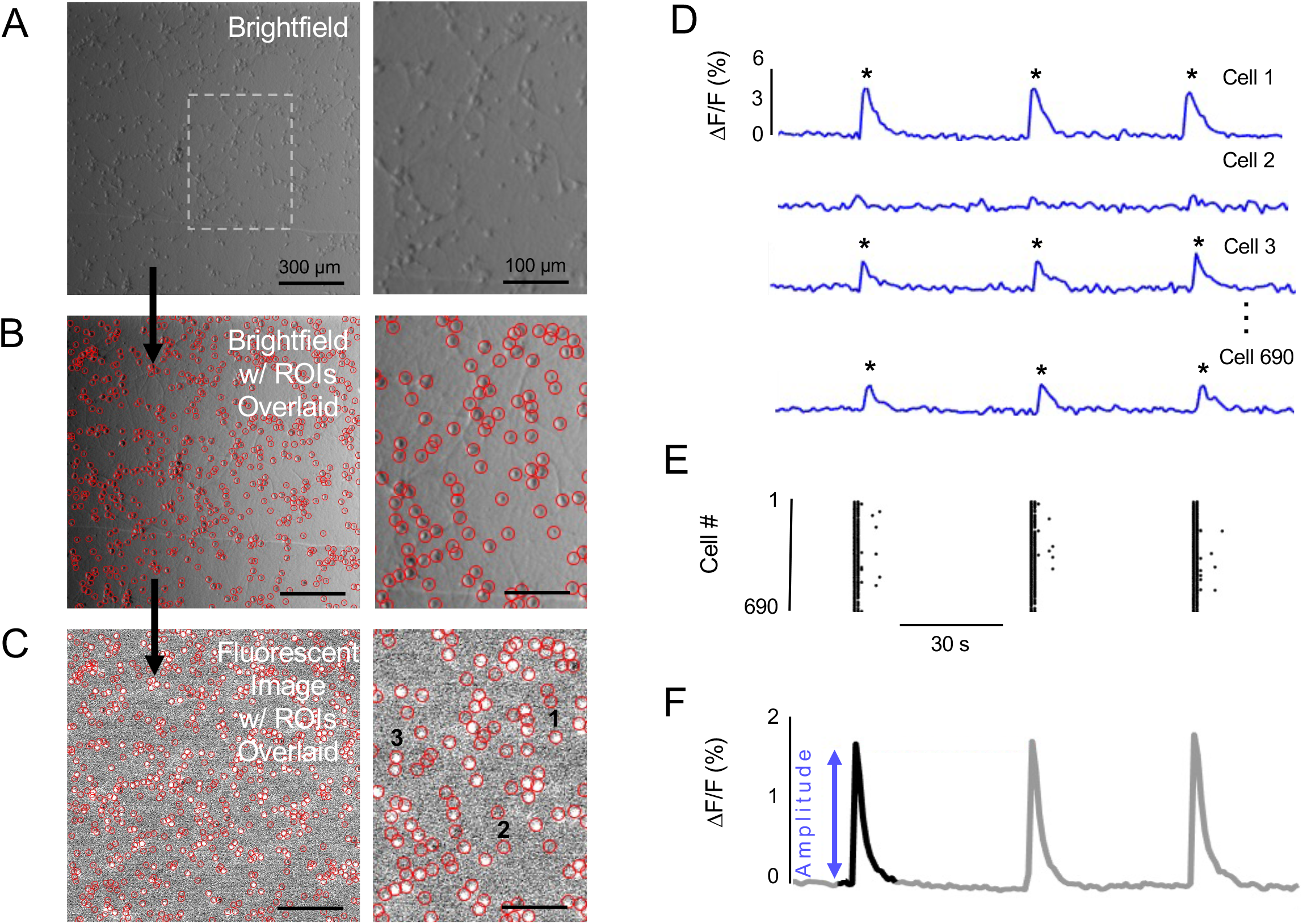
Ca^2+^ activity of individual cells. (A) Brightfield image of 690 cells at 4x magnification; (right) enlarged region defined by dashed rectangle. (B) Brightfield image of the same field in (A) with circular ROIs overlaid; (right) enlarged image showing alignment of ROIs with cell bodies. (C) ∆F/F image of the same field in (A) with ROIs overlaid; (right) enlarged image to show that ROIs also coincide with active (bright) cells and non-active cells. (D) Example individual Ca^2+^ signals shown as ∆F/F for active and non-active cells. Detected Ca^2+^ events are shown with an asterisk (*). (E) A raster plot of cell Ca^2+^ event detections across time. (F) Fluorescence intensity measured over the FOV during the same time period in which fluorescence from individual ROIs were measured.

Accuracy for the positioning of ROIs over neuronal somata was determined using five culture wells. Across these samples, the false positive rate was 1.6±0.2% (Mean ± S.E.M.) and the false negative rate was 4.6±1.7%. The low false positive and false negative rates demonstrated high accuracy of the macro without manual interventions.

Detection of Ca^2+^ activity from individual cell ∆F/F values (Figure 1D) relied on a filtering and numerical threshold scheme written in MATLAB. Baseline drift in fluorescence was removed with a Savitzky-Golay filter function. For event detection, a numerical threshold was set to greater than 3-standard deviations above the mean baseline fluorescence signal. The time for the peak ∆F/F of each detected event was recorded for every cell in the image stack.

The results of this event detection procedure for each image stack were used to generate a raster plot of cellular activity against time (Figure 1E). These data were also used to calculate the percentage of cells participating during periods of synchronous activity (participation rate). The raster plots of individual neuronal activity were highly correlated with the recordings of fluorescence intensity averaged over the entire FOV (Figure 1F). This correspondence between individual neuronal activity and synchronous activity across the population of neurons indicated that the neuronal networks have a high degree of functional connectivity, as previously reported (Kamioka et al., 1996; van Pelt et al., 2004; Patel et al., 2012).

### 2.2. Mapping Connections with LSPS

Neurons were selected for mapping of network connections if they participated in a synchronous Ca^2+^ event. For each mapping session, 75 active neurons closest to the center of the FOV were selected as targets. The soma of each selected neuron (black circle in Figure 2A-B) was photostimulated (Nguyen et al., 2017) and the corresponding Ca^2+^ responses of the target neuron (black ∆F/F trace in Figure 2B) and other neurons in the FOV (red circles and traces in Figure 2B) were recorded for later analysis.

**Figure 2.**
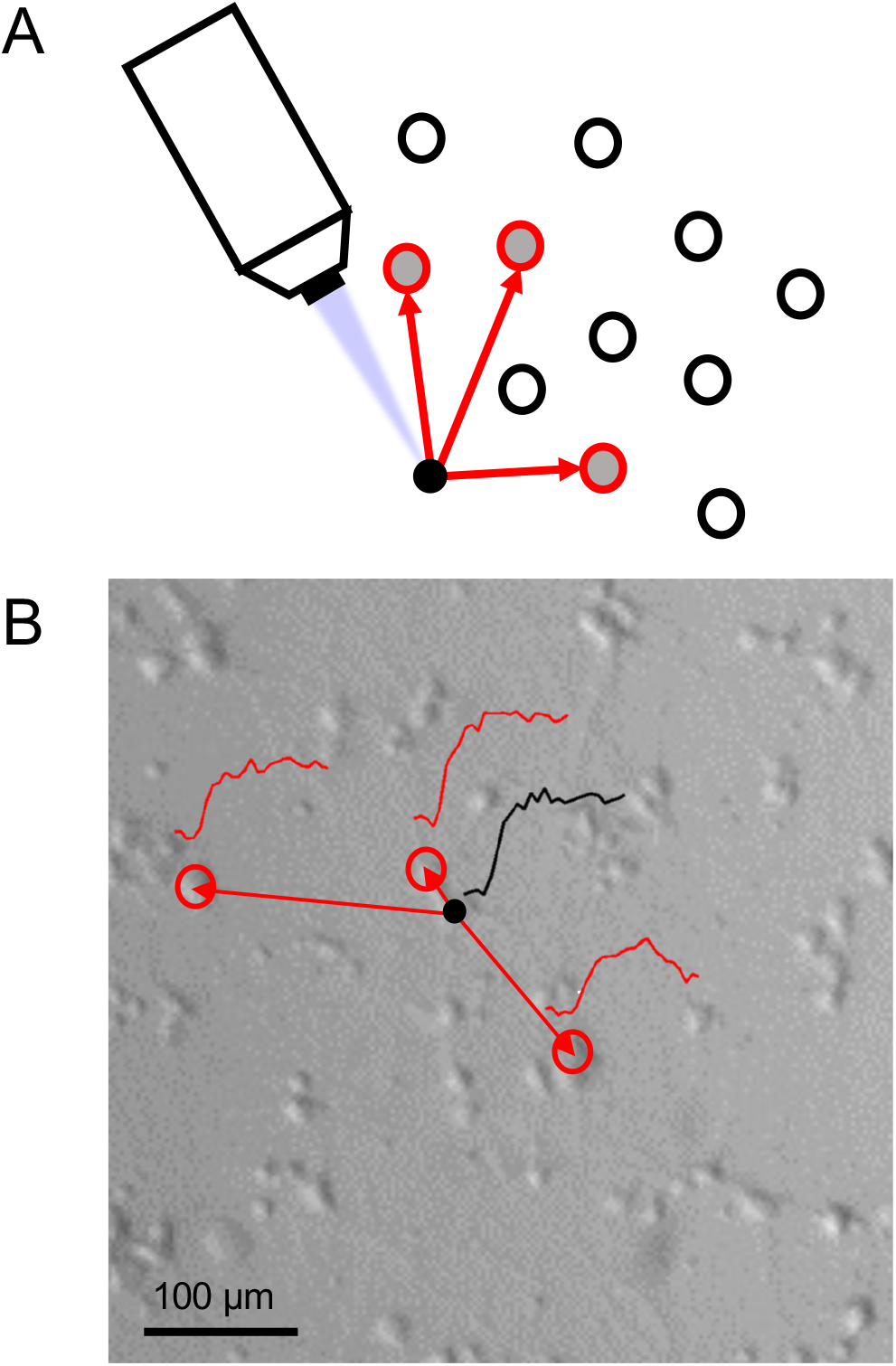
Mapping with LSPS technique. (A) Diagram of the LSPS technique: UV laser light photolyzes caged glutamate surrounding the target cell (●). Subsequent activity of target cell and other cells (o) responding to this stimulation are recorded as increases in Ca^2+^ fluorescence. A total of 75 neurons can be stimulated within 10 minutes. (B) A representative connectivity map of one neuron is shown along with LSPS-induced Ca^2+^ responses against a brightfield image with cells.

When mapping was repeated for a second session (as shown in Figures 5,6,8-11), the target neurons were photostimulated in the same order. Furthermore, photostimulation was avoided during synchronous Ca^2+^ activity events in order to not misinterpret spontaneous activation as a stimulated response. If synchronized Ca^2+^ activity occurred during photostimulation, mapping was stopped and reinitiated after the event.

A MATLAB routine generated a series of a potential links/connections between targeted neurons and neurons exhibiting Ca^2+^ responses after each photostimulation. The criteria for designating a potential link were based on the amplitude and timing of the Ca^2+^ responses. First, the amplitude of Ca^2+^ response of the responding neuron had to exceed the 3 standard deviation threshold value defined previously. Second, the Ca^2+^ response had to begin within a window of 300 ms or less after photostimulation of the targeted neuron. This delay window is based on the finding of Peterlin et al. (2000) that large increases of Ca^2+^ in a neuron monosynaptically connected to a patch-clamped neuron typically occurred about 300 ms after the first action potential in the patch-clamped neuron. A longer delay between LSPS and a Ca^2+^ response might indicate a polysynaptic connection or spontaneous activity unrelated to functional links between neurons. Third, the response had to exceed the threshold value for at least 1 second. Ca^2+^ responses in neurons had to meet these criteria to be accepted as a response linked to the photostimulated neuron (Figure 3A). Increases in ∆F/F that were accepted as connections were individually confirmed by manually checking image stacks in a MATLAB GUI. Ca^2+^ responses that did not meet these criteria were rejected. As examples, the other recordings of ∆F/F shown in Figure 3 were rejected because they resulted from random noise in the ∆F/F signal (Figure 3B) or began more than 300 ms after photostimulation (Figure 3B and C).

**Figure 3.**
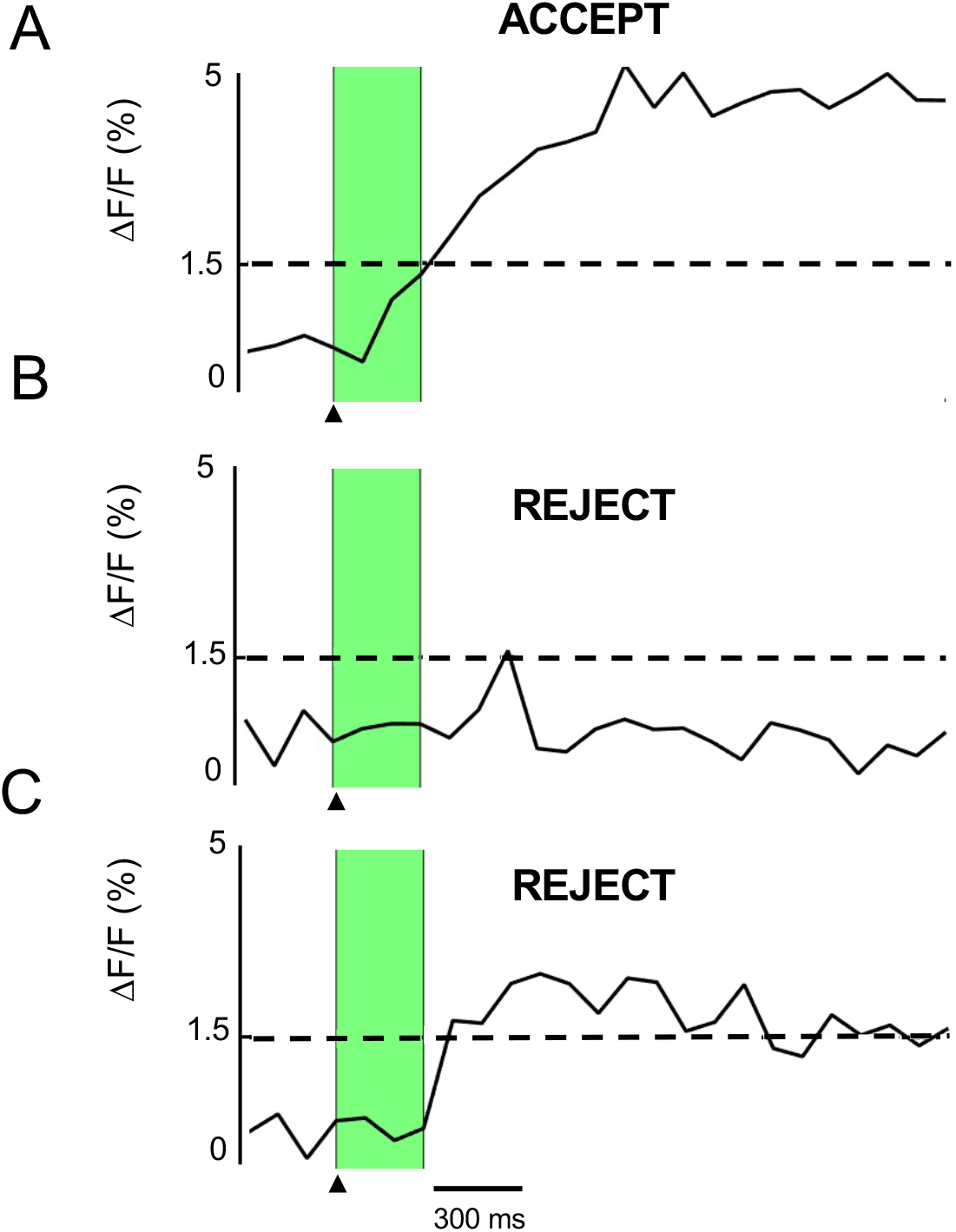
Connection criteria. ▲ indicates when photostimulation occurs. The green bar indicates the 300 ms window after photostimulation during which ΔF/F must exceed the threshold value (dashed line) to be accepted as a link. (A) An accepted link based on the acceptance criteria. (B) A response that met the ΔF/F threshold but rejected as noise. (C) A rejected link that met the threshold but occurred after the accepted window (> 300ms).

For each session, a connectivity map of directional links, i.e. from target cell to responding cell, was generated from this analysis and overlaid on top of the bright field image (Figure 2B). These maps provided a spatial view of the network and its connections.

### 2.3. Excitatory-only neuronal networks (ENNs) demonstrate stable parameters with time

To generate networks involving only excitatory neurotransmission, bicuculline (40 µM) was added to solutions to block inhibitory GABAergic neurotransmission. Consistent with prior literature, bicuculline reduced frequency and increased the amplitude of synchronous Ca^2+^ activity (Goforth et al., 2011). ENNs were used for all subsequent experiments.

Fluorescence imaging of ENNs showed stable activity with time (Figure 4A) as evidenced by the relatively conserved amplitude and frequency of synchronous Ca^2+^ elevations during recordings periods up to 1 hour in duration. In order to effectively combine both fluorescence imaging of synchronous Ca^2+^ activity and mapping with LSPS, the experimental paradigm in Figure 4B was established. This experimental paradigm enabled comparisons of various network characteristics (extracted from connectivity maps) before and after specific experimental interventions, eg. application of pharmacological agents.

**Figure 4.**
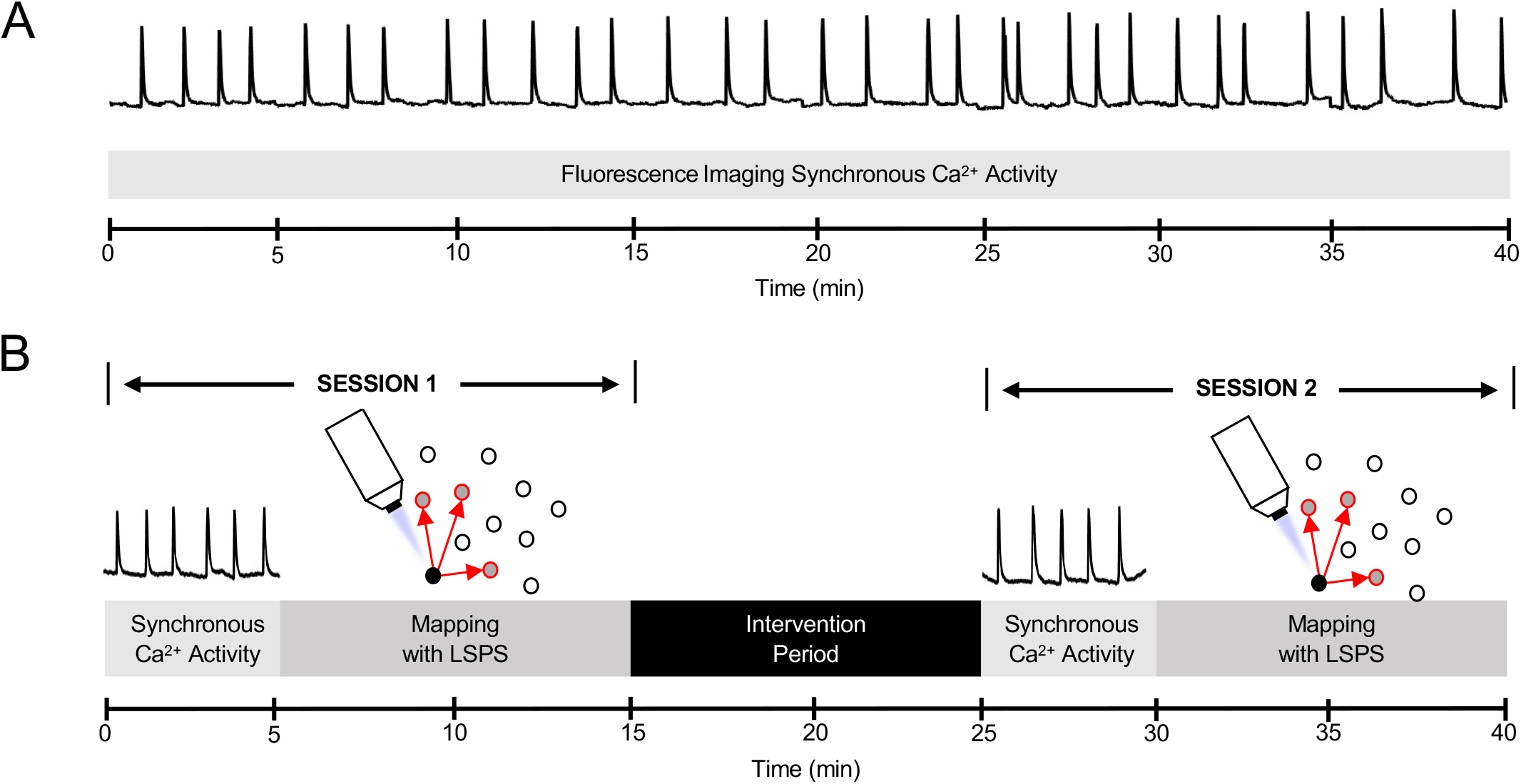
Experimental paradigm. (A) Synchronous Ca^2+^ activity over time demonstrating stability of amplitude and frequency in the absence of any interventions or LSPS mapping. (B) The experimental paradigm combining fluorescence imaging of synchronous Ca^2+^ activity and mapping with LSPS before and after an intervention period.

**Figure 5.**
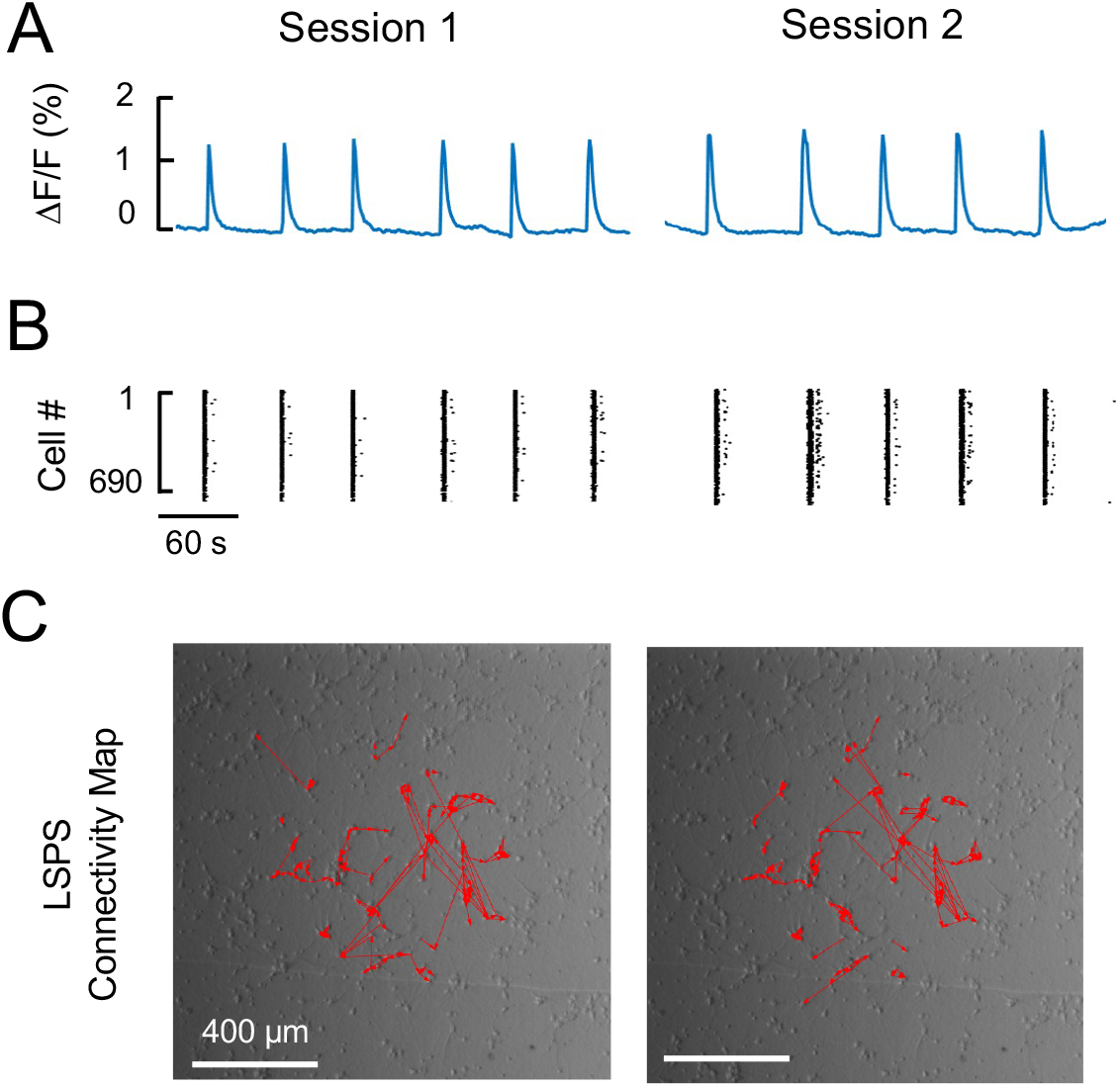
Effects of control conditions on parameters with time. A representative experiment is shown for a control experiment without any interventions. (A) Synchronous Ca^2+^ activity. Mean amplitude was 1.68% and mean frequency was 20.2 mHz in Session 1 and 1.93% and 18.6 mHz, respectively, in Session 2. (B) Raster plot of spontaneous activity. Participation rate was 79.1% and 87.4% in Sessions 1 and 2, respectively. (C) LSPS mapping of 75 neurons. In Session 1, 72 active neurons had a total of 167 links. The *k* value = 2.3, and *C* = 0.25. In Session 2, 71 neurons remained active with 171 links, with *k* = 2.4 and *C* = 0.25. Comparing links between LSPS sessions, total links increased 2.4%, 75.4% of links remained stable, 24.6% of links were lost and 26.9% of links were new.

**Figure 6.**
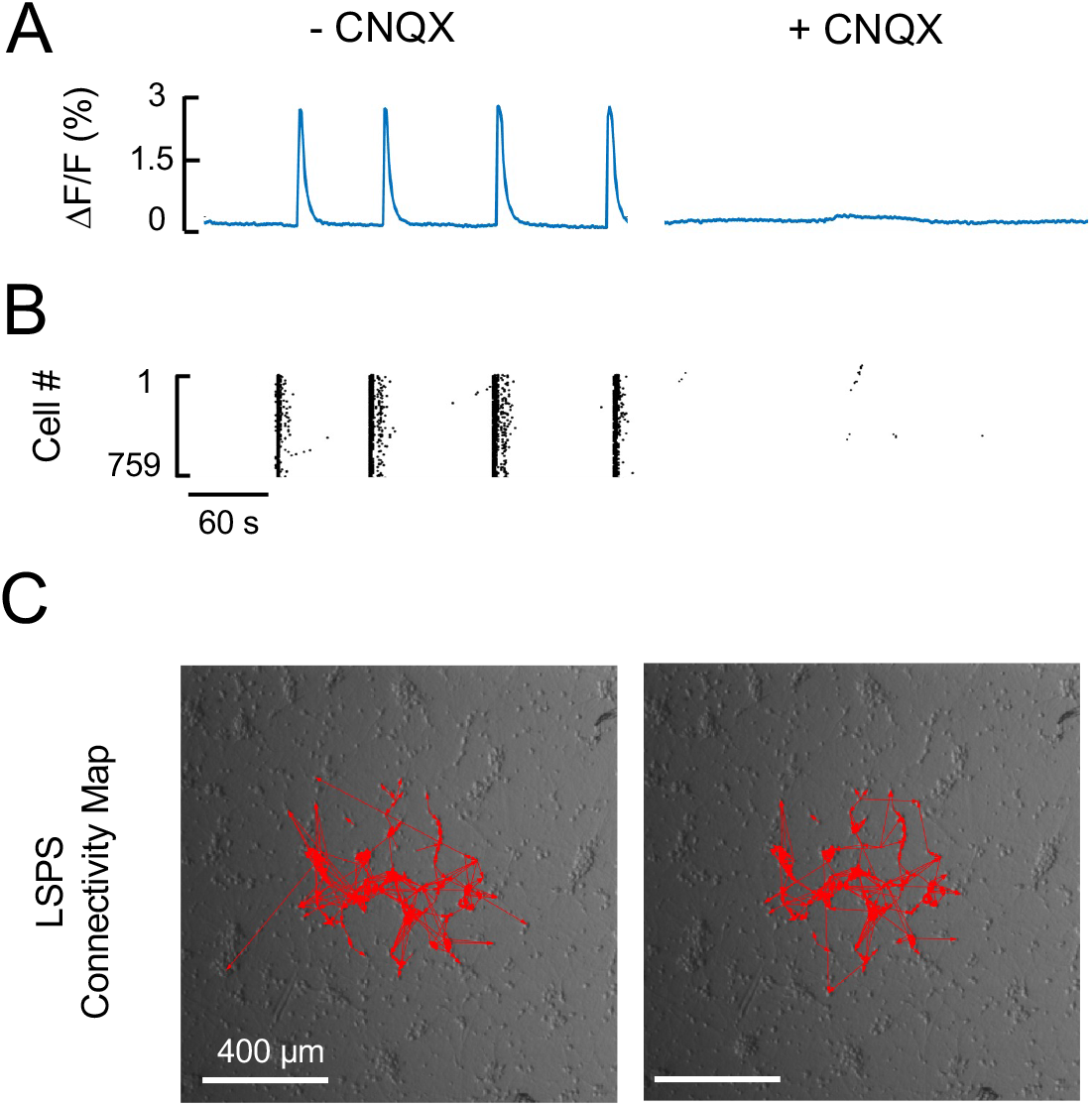
Effects of CNQX on synchronous activity and LSPS mapping. (A) Synchronous Ca^2+^ activity. Mean amplitude was 2.74% with a mean frequency of 13.9 mHz before CNQX. After 10 minutes in the presence of CNQX, synchronous Ca^2+^ activity was abolished. (B) Raster plot of synchronous activity. The participation rate of 97.6% prior to CNQX addition was abolished after 10 minutes in the presence of CNQX. (C) LSPS mapping of 75 neurons. Prior of CNQX addition, 68 neurons were active with a total of 365 links. *k* = 5.4 and *C* = 0.35. After 10 minutes in the presence of CNQX, 68 neurons were active with 332 links. *k* = 4.9 and *C* = 0.42. Comparing links between sessions, total links decreased 9%, 80.0% of links were stable, 20.0% of links were lost and 11.0% of links were new.

Three parameters were determined for events marked by synchronous Ca^2+^ activity. Event amplitude was defined as the peak ∆F/F signal averaged over the entire FOV. Event frequency was defined as the number of synchronous Ca^2+^ activity events per second. Both of these parameters are similar to those described in previous work (Patel et al., 2014). Participation rate was defined as the percentage of cells participating in synchronous Ca^2+^ activity.

Results of a representative control experiment are shown in Figure 5. Synchronous Ca^2+^ activity (Figure 5A) and raster plots (Figure 5B) appeared to be largely unchanged before and after the intervention period. Synchronous Ca^2+^ activity had an average event amplitude of 1.68% and a frequency of 20.2 mHz (Figure 5A, Session 1). After the 10 min intervention period, in which no intervention occurred, synchronous Ca^2+^ activity had an average amplitude of 1.93% and a frequency of 18.6 mHz (Figure 5A, Session 2). Raster plot activity demonstrated a participation rate before and after the intervention period 78.8% and 86.7%, respectively (Figure 5B).

In accordance with the experimental paradigm (Figure 4), LSPS mapping was performed prior to and after the intervention period, which in this case had no intervention. The connectivity maps resulting from LSPS of the same 75 neurons in each session before and after the intervention period are shown in Figure 5C. Similar to the synchronous Ca^2+^ activity, the connectivity maps looked largely unchanged.

To quantify connectivity between neurons, two parameters were determined. The number of responsive neurons was defined as number of neurons, out of the 75 targeted by LSPS, that elicited a Ca^2+^ response in the target cell (see Figure 2). The total number of links (# Links) was defined as the number of linked Ca^2+^ events (see Figure 2 and 3) occurring in non-targeted neurons after LSPS of the 75 targeted neurons. In Figure 5C, the number of active neurons was unchanged in the two mapping sessions. The # of links, 167 before and 171 after the intervention period, were also similar. Thus, as suggested by the connectivity maps, the overall connectivity parameters in this experiment were largely unchanged in the absence of any intervention.

LSPS mapping also allows for the properties of a functioning neuronal network to be defined. In our analysis, we used a basic measure of network connectivity, average degree (*k*) and a measure of segregation, average clustering coefficient (*C*), to describe the properties of these networks. Average degree, effectively the average number of links per neuron, is defined as the # links divided by the number of LSPS targeted neurons that produced a Ca^2+^ response. In Figure 5C, *k* was calculated to be 2.3 and 2.4 before and after the intervention period, respectively. Average clustering coefficient quantifies the extent to which neurons and their neighbors are interconnected to each other, where the clustering coefficient for each neuron is the fraction of the maximum number of possible connections between all the neurons linked to that neuron (Rubinov and Sporn, 2010). In this experiment, *C* was 0.25 both before and after the intervention period. As suggested by Figure 5C, these network parameters are stable in this control experiment.

To understand the properties of links identified during the mapping sessions with LSPS, links were classified into 3 categories, stable, lost and new. Stable links were those present in both mapping sessions before and after the intervention period. Lost links were those that were present before the intervention period, but absent after. New links were those that were absent before the intervention period, but present after. In the experiment shown in Figure 5, 75.4% of links were stable, 24.6% were lost, and 26.9% were new. Similar to network connectivity parameters, the high percentage of stable links indicated that, under control conditions, connectivity between neurons was consistent, i.e. lost links were essentially balanced by the appearance of new links.

In addition to categorizing neuronal links, the properties of stable links could also be characterized with two additional parameters: change in glutamate responsiveness and weight of stable links. Change in glutamate responsiveness was determined by comparing the amplitudes of the Ca^2+^ responses for each targeted neuron before and after the intervention period. In Figure 5, the amplitudes of the Ca^2+^ responses of the 75 targeted neurons slightly increased between the LSPS mapping sessions (9.0 ± 5.7%; Mean ± S.E.M.). Change in the weight of stable links was determined by comparing the amplitudes of the Ca^2+^ responses for each responding neuron before and after the intervention period. The amplitudes of the Ca^2+^ responses of the 179 linked neurons in Figure 5 were also increased (13.8 ± 2.7%; Mean ± S.E.M.). These data show that, over the time course of this experiment, the response to LSPS became somewhat larger in targeted and responding neurons connected by stable links.

Parameters derived from synchronous Ca^2+^ activity and LSPS were calculated for ten control experiments. Two of the parameters for synchronous Ca^2+^ events, amplitude and participation rate, demonstrated small, but significant increases, while frequency displayed a small, but significant decrease (Table 1). LSPS parameters, # links *k* and *C*, also had small but statistically significant increases, while the number of responsive neurons was unchanged (Table 1). Thus, in general, the network became slightly more connected over the time course of control experiments. Analysis of the link properties showed that 81.3% [9.8%] (median [IQR]) of links were stable, 18.7% [9.8%] were lost, and 30.7% [20.1%] were new. Thus, the slight increase in network connectivity resulted from the appearance of new links relative to the number of lost links. Stable links also showed small but significant increases in glutamate responsiveness (5.8 ± 3.6%; Mean ± SEM) and weight of stable links (7.2 ± 1.0%). Altogether, these data suggest that the combined techniques of fluorescence imaging of synchronous Ca^2+^ activity and mapping with LSPS can provide a detailed description of the properties of neuronal networks. Furthermore, over the time course of our control experimental protocol, these parameters are either unchanged or show only small changes.

**Table 1:** Network Parameter Values. Data are displayed as median with its interquartile range (IQR) in brackets. Replicates (n = 10) were performed in 3 cultures from 3 animals. * p < 0.05 using Wilcoxon matched pairs signed rank tests.

| Parameter | Control<br>(No Intervention) |  |
| --- | --- | --- |
|  | Session 1 | Session 2 |
| *Parameters from Synchronous $\text{Ca}^{2+}$ Activity* |  |  |
| **Amplitude**<br>($\Delta F/F$) (%) | 1.3<br>[0.8] | 1.4<br>[1.2] * |
| **Frequency**<br>(mHz) | 17 [12] | 14 [9] * |
| **Participation Rate (%)** | 64 [24] | 67 [30] * |
| *Parameters from LSPS Mapping* |  |  |
| **# Links** | 150 [66] | 168 [81] * |
| **Responsive Neurons** | 73 [9] | 71 [7] |
| **$k$** | 2.2 [1.0] | 2.4 [1.0] * |
| **$C$** | 0.20 [0.10] | 0.25 [0.12] * |

### 2.4. AMPA receptor blockade does not affect LSPS mapping-derived parameters

Excitatory transmission in cortical networks are mediated mainly by AMPA and NMDA receptors (Nusser, 2000). To evaluate the effect of AMPA receptor blockade on network parameters, 6-cyano-7-nitroquinoxaline-2,3-dione (CNQX, 2 µM) was added to the bath solution at the beginning of the 10-minute intervention period. As shown in Figure 6A, AMPA-blockade resulted in a complete suppression of synchronous Ca^2+^ activity, and thus no participation in synchronous events (Figure 6B). Loss of spontaneous activity in the presence of CNQX has previously been reported (Murphy et al., 1992; Suresh et al., 2016).

In contrast, the map of LSPS links was largely unaffected (Figure 6C) with the number of responsive neurons being unchanged and the total number of links decreasing by 9%. Network parameters were 5.4 and 4.9 for *k* before and after CNQX, respectively, and 0.35 and 0.42 for *C*, respectively. Stable, lost and new links were 80.0%, 20.0% and 11.0% of total links, respectively. Thus, in this experiment, CNQX had only small effects on LSPS derived parameters.

Overall, complete suppression of synchronous Ca^2+^ activity by CNQX was observed in four out of five experiments. When synchronous Ca^2+^ activity was not completely suppressed, amplitude and participation rate appeared unchanged, but frequency was dramatically decreased (not shown).

By comparison, LSPS mapping-derived connectivity (Figure 7A) and network parameters (Figure 7B) in these five experiments were not significantly different before and after CNQX, and similar to those in control experiments. In addition, the percentages of stable, lost and new links were not significantly different than in control experiments (Figure 7C). Overall, for these five experiments, the glutamate responsiveness (21.4 ± 2.7 %; Mean ± S.E.M.) and weight of the stable links displayed relatively small, but significant increases in the presence of CNQX (9.0 ± 1.0%; Mean ± S.E.M.), similar to control conditions. These results show that spontaneous network activity in mixed neuronal cultures is largely dependent on AMPA glutamatergic synaptic transmission. In contrast, network connections triggered by localized photolysis of caged glutamate at a single neuron were largely independent of AMPA receptor-dependent neurotransmission.

**Figure 7.**
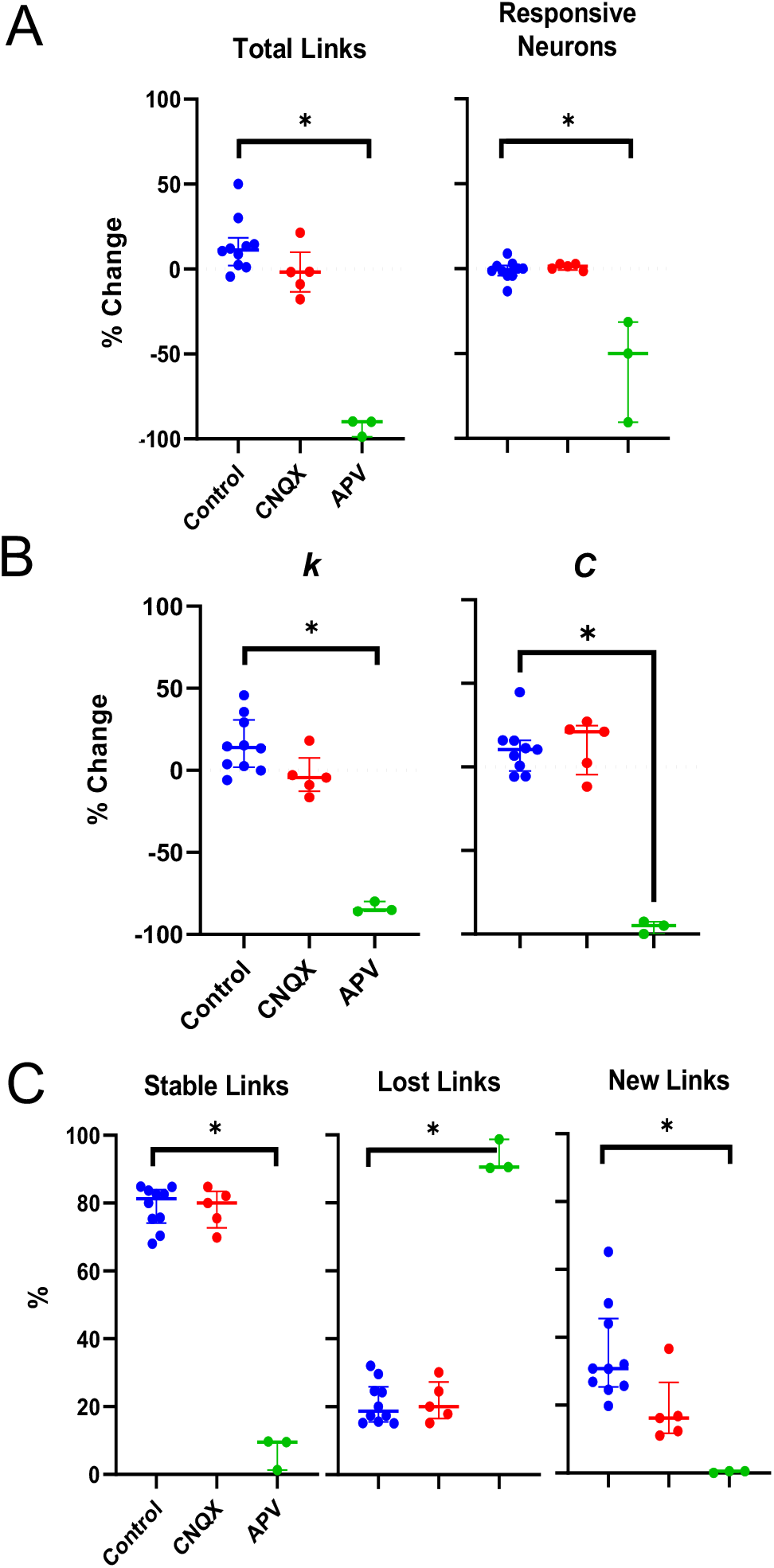
Changes of LSPS mapping derived network parameters after glutamate receptor blockade. Individual data points are shown with the median and the interquartile range (25% to 75% IQR). Data are displayed as percent change for mapping (A) and network parameters (B), and as percent for link properties (C) for control, CNQX and APV experiments. * p < 0.05 using Kruskal-Wallis with post-hoc Dunn’s correction comparing groups. n = 10 for control, n = 5 for CNQX and n = 3 for APV. For CNQX experiments, total links (median [IQR]) were 173 [146] and 176 [125], and the number of responsive neurons was 72 [3] and 74 [5], before and after drug addition, respectively. For APV experiments, total links (median [IQR]) were 297 [232] and 20 [40], and the number of responsive neurons were 73 [7] and 33 [43], before and after drug addition, respectively.

**Figure 8.**
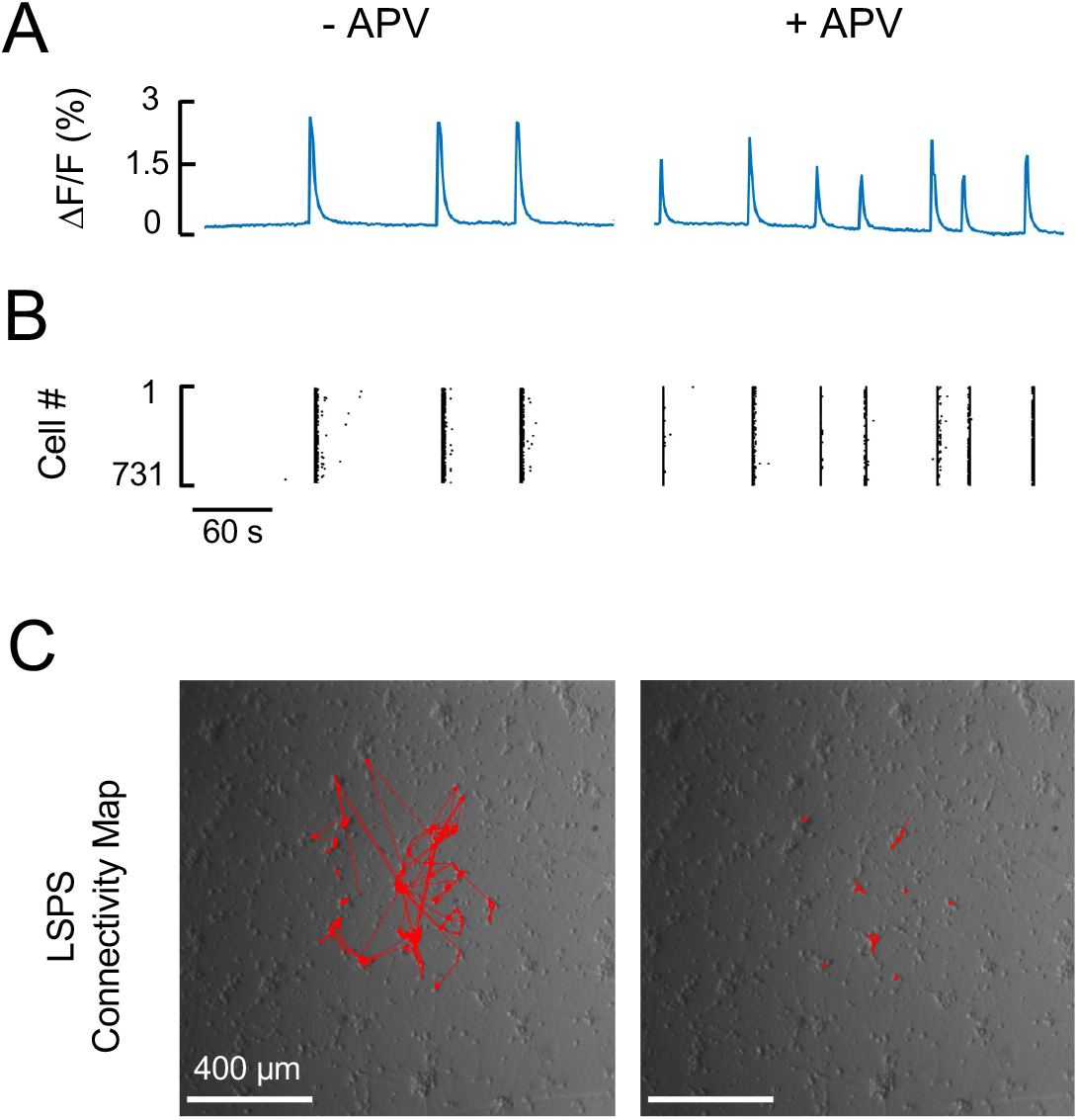
Effects of APV on synchroous activity and LSPS mapping. (A) Synchronous Ca^2+^ activity. The mean amplitude of 2.49% with a mean frequency of 13.6 mHz before APV addition. After 10 minutes in the presence of APV, mean amplitude was 1.64% with a mean frequency of 23 mHz. (B) Raster plot of synchronous activity. Participation rate of 94.8% prior to APV addition, and after 10 minutes in the presence of APV, participation rate was 71.6%. (C) LSPS mapping of 75 neurons. Prior to addition of APV, 66 neurons were active with a total of 199 links. *k* = 3.0 and *C* = 0.27. After 10 minutes in the presence of APV, 33 neurons were active with a total of 20 links. *k* = 0.6 and *C* = 0.01. Comparing between sessions, total links decreased 89.9%, 9.5% of links remained stable, 90.5% of links were lost and 0.5% of links were new.

### 2.5. NMDA receptor blockade suppresses LSPS linkages

To evaluate the dependence of network parameters on NMDA-dependent neurotransmission, D-2-amino-5-phosphono-valerate (APV, 25 µM) was added to the bathing solution at the beginning of the 10-minute intervention period. A representative experiment in Figure 8 A and B shows that NMDA-blockade resulted in an overall decrease in amplitude of and participation rate in synchronous Ca^2+^ activity (not shown), though with greater variability. Frequency of events, on the other hand, was increased. Before APV addition, the amplitude, frequency and participation rate were 2.49%, 13.6 mHz, and 94.2%, respectively. After APV addition, the amplitude, frequency and participation rate were 1.64%, 23 mHz, and 71.6%, respectively.

LSPS derived parameters were also affected by addition of APV. The number of responsive neurons was decreased by 50% after APV addition. Network connectivity maps (Figure C) showed that APV markedly reduced links between neurons. Network parameters, *k* and *C*, were also decreased. Only 9.5% of links were stable, while 90.5% were lost and only 0.5% of links were new. Thus, APV decreased the stability of links and reduced the likelihood of new links, thereby greatly inhibiting LSPS triggered connections between neurons.

Similar results were observed in two additional experiments. APV dramatically reduced the number of links and responsive neurons (Figure 7A). Likewise, network parameters *k* and *C* were also statistically decreased (Figure 7B), though the latter parameter might be biased by the lack of connectivity in the presence of APV (Rubinow and Sporns, 2010). The percentage of stable links was also greatly reduced, accounting for the large increase in lost links (Figure 7C). In addition, the percentage of new links was significantly decreased. Overall, for these three experiments, glutamate responsiveness of targeted neurons was decreased by 63.6 ± 1.6%, and the weight of stable links for responding neurons was decreased by 65.0 ± 1.4% (Mean ± SEM). Though both of these parameters were reduced by APV, a linear regression analysis showed that the change in the weight of stable links was only very weakly correlated with the change glutamate responsiveness (R^2^ = 0.11). Altogether, these data show that APV dramatically suppressed links probed by LSPS and decreased strength of the remaining links.

To confirm that NMDA receptor dependent neurotransmission was critical for neuronal links identified by LSPS, the effect of increasing extracellular Mg^2+^ on link properties was examined. Increasing extracellular Mg^2+^ from 1 mM to 2 mM during the intervention period significantly reduced the number of observed links (Figure 9) with little effect on the number of responsive neurons, as would be expected if Mg^2+^ inhibits NMDA receptor dependent neurotransmission (Nowak et al., 1984). Similar results were observed in a second experiment.

**Figure 9.**
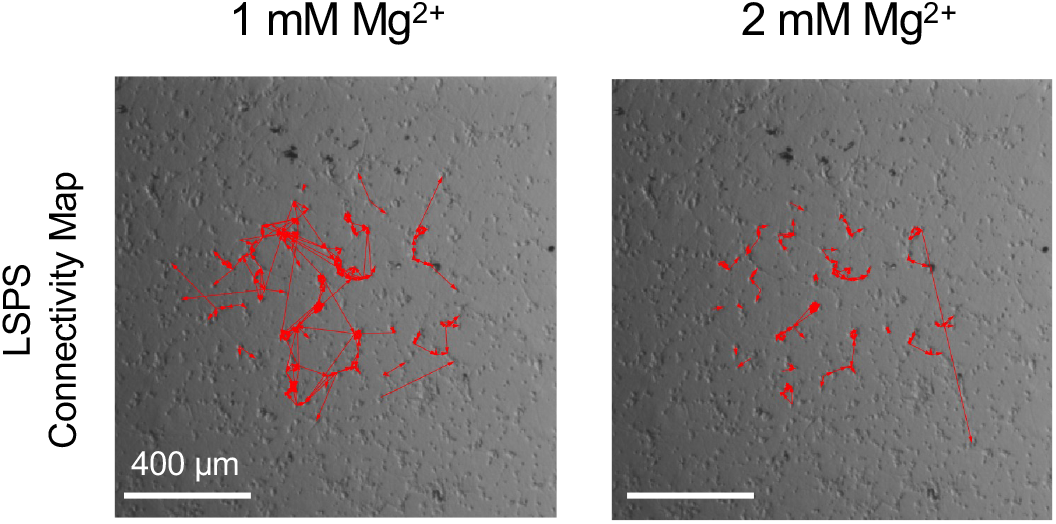
Effect of increased extracellular Mg^2+^ on LSPS connectivity maps. At 1 mM Mg^2+^, there were 228 links with 75 responsive neurons. At 2 mM Mg^2+^, links were reduced to 116 with 72 responsive neurons. Increasing Mg^2+^ resulted in 49.6% of links remaining stable, 50.4% lost and 1.3% new.

Together with data showing a lack of effect of CNQX, LSPS-derived network parameters are clearly dependent on NMDA receptor-dependent neurotransmission. Thus, LSPS provides are unique window into understanding the role of NMDA-dependent neurotransmission properties that is not available by monitoring synchronous Ca^2+^ activity.

### 2.6. Modulation of network properties by manipulating GluN2A receptor-dependent neurotransmission

To further illustrate the utility of the LSPS approach for investigating changes in network properties, the effect of NMDA subtype receptor manipulation was tested by examining the effect of pharmacological inhibition and positive modulation of GluN2A receptor-dependent neurotransmission on synchronous Ca^2+^ activity and neuronal links determined by LSPS mapping. GluN2A specific receptor blockade was produced by addition of 50 nM NVP-AAM077 (Massey et al., 2004) during the intervention period. Receptor blockade produced, on average, a 28% decrease in the amplitude of synchronous Ca^2+^ activity (Figure 10A). In addition, the participation rate in synchronous events decreased from 77% to 52%. In contrast, the frequency of spontaneous activity was increased by 133%. Similar results were observed in two other experiments.

**Figure 10.**
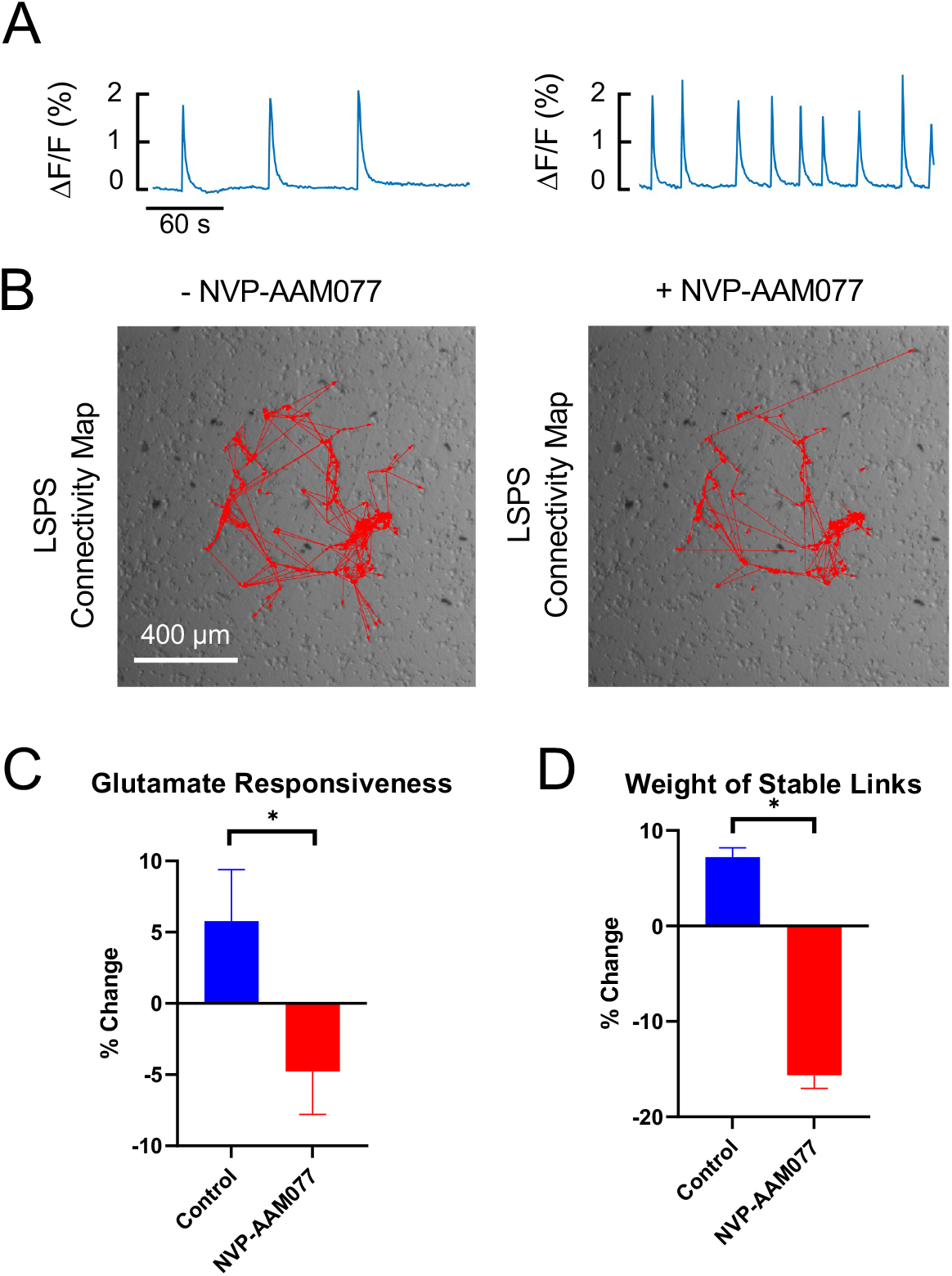
Effect of a GluN2A blocker on synchronous activity and LSPS Mapping. (A) Synchronous Ca^2+^ activity. Mean amplitude and frequency were of 2.01% and 16.9 mHz, respectively, before and 1.45% and 39.3 mHz after 10 minutes in the presence of 50 nM NVP-AAM077. (B). LSPS connectivity map before and after addition of NVP-AAM077. (C) Percent change in glutamate responsiveness for experiments without NVP-AAM077 (n = 750 neurons) and with NVP-AAM077 (n = 225 neurons). (D) Percent change in weight of stable links without NVP-AAM077 (n = 1196 links) and with NVP-AAM077 (n = 484 links). Paired t-tests indicated significant differences between groups (*p < 0.05). Data displayed as Mean ± S.E.M. Total n = 10 wells without NVP-AAM077, and n = 3 wells with NVP-AAM077.

As a converse experiment, GluN2A receptor function was enhanced by adding the subtype specific positive modulator, GNE-0723 (Hanson et al., 2020), during the intervention period. For these experiments, the intervention period was increased to 20 minutes duration to provide enough time for drug action to occur. Control experiments demonstrated that network parameters were not different whether the intervention period was 10 or 20 minutes in duration (not shown). GNE-0723 had little effect on the average amplitude of synchronous Ca^2+^ events (Figure 11A). Likewise, participation rate, 92.7% before and 95.0% after drug addition, was unaffected. The frequency of synchronous events, however, did increase by 77%. Similar results were observed in a second experiment.

**Figure 11.**
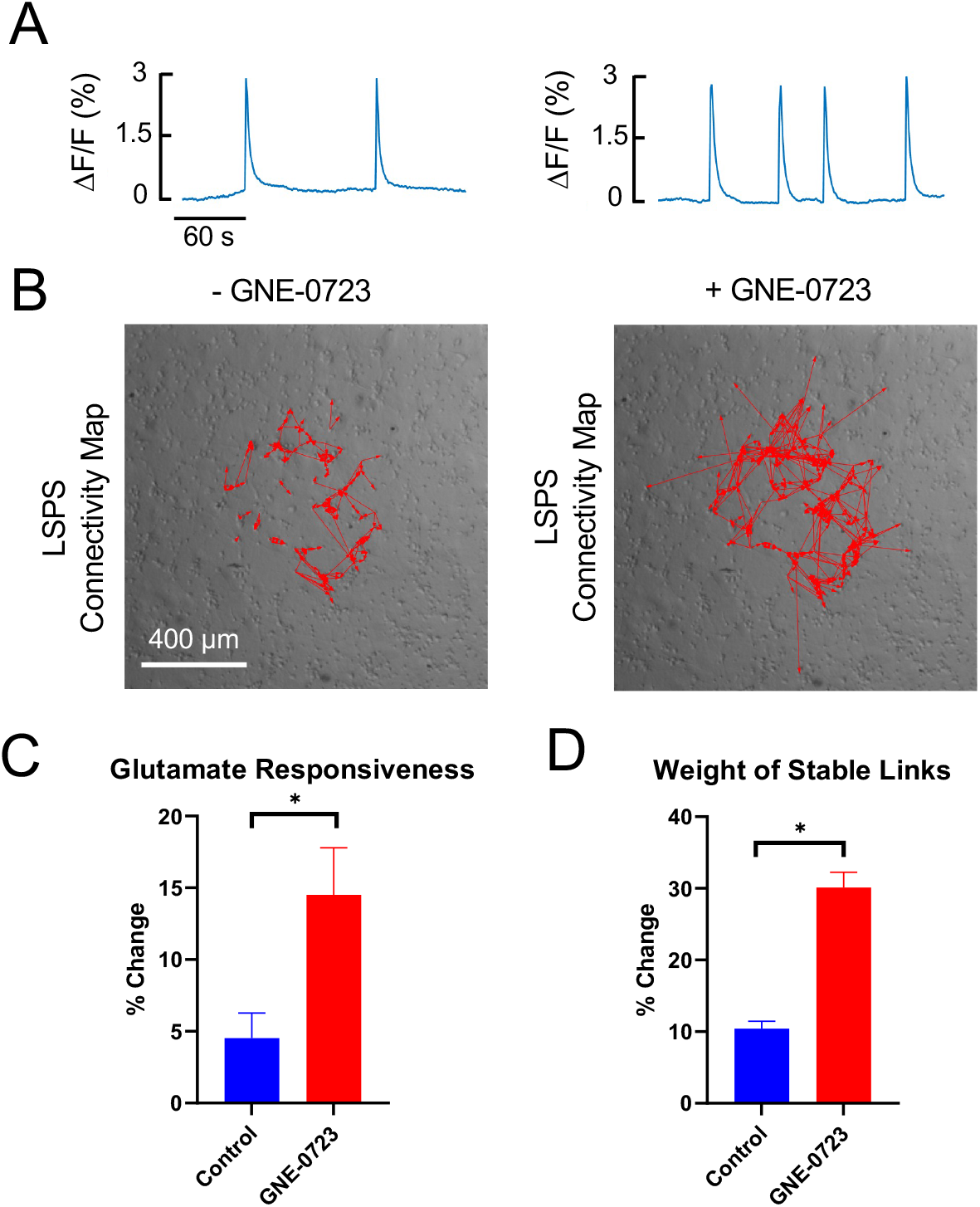
Effect of a GluN2A positive modulator on spontaneous activity and LSPS mapping. (A) Synchronous Ca^2+^ activity. Mean amplitude and frequency were of 2.83% and 9.1 mHz, respectively, before and 2.92% and 16.1 mHz before and after 20 minutes in the presence of 1 µM GNE-0723 (B). LSPS connectivity map before and after addition of 1 µM GNE-0723. (C) Percent change in glutamate responsiveness of experiments without GNE-0723 (n = 450 neurons) and with GNE-0723 (n = 150 neurons). (D) Percent change in weight of stable links without GNE-0723 (n = 1168 links) and with GNE-0723 (n = 529 links). Paired t-tests indicated significant differences between groups (*p < 0.05). Data displayed as Mean ± S.E.M. Total n = 6 wells without GNE-0723 and n=2 wells with GNE-0723.

Though pharmacological interventions used in these experiments are predicted to have opposite effects on GluN2A-dependent neurotransmission, the effects on synchronous Ca^2+^ events were not always qualitatively different. These results suggest that interpretation of the effects of manipulating neuronal transmission on network functionality is complicated when using parameters derived from spontaneous activity and can critically depend on which parameters are compared.

The effect of GluN2A blockade on links determined by LSPS is shown in Figure 10B for the same experiment shown in Figure 10A. Prior to addition of NVP-AAM077, 408 links were identified in 75 responsive neurons, out of the 75 neurons tested, with a *k* of 5.4 and C of 0.38. After drug addition, the number of responsive neurons was unchanged, 75 out of 75 tested, but the number of links was reduced to 251, *k* was reduced to 3.3 and *C* decreased to 0.30. Overall, 56.1% of the links were stable in the presence of NVP-AAM077, 43.9% were lost and 5.4% were new. These values are much different than those observed in control experiments (Figures 5 and 7). Similar results were observed in all 3 experiments in which GluN2A blockade was tested. In addition, glutamate responsiveness and weight of stable links was also decreased by drug application (Figure 10C and D, respectively). These data demonstrated that GluN2A receptor inhibition decreased links between neurons and reduced the strength of stable links.

The effect of GluN2A positive modulation on links determined by LSPS is shown in Figure 11B for the same experiment shown in Figure 11A. Prior to addition of GNE-0723, 200 links were identified in 74 responsive neurons, out of the 75 neurons tested. The *k* value was 2.7 and *C* was 0.28. After drug addition, the number of responsive neurons was essentially unchanged, 75 out of 75 tested, but the number of links increased to 453, *k* increased to 6.0, and the C also increased to 0.37. Overall, 89.5% of the links were stable in the presence of GNE-0723, 10.5% were lost and 137% were new. Similar results were observed in another experiment. Thus, positive modulation increased the number of links between neurons, largely by adding new functional links between neurons and by increasing connectivity between active neurons. In addition, GNE-0723 increased glutamate responsiveness and weight of stable links between neurons (Figure 11C and D, respectively). Thus, positive modulation of GluN2A strengthened existing links between neurons and promoted activity in other links that otherwise were inactive.

Altogether, these data show that LSPS provided a consistent and detailed view of how manipulating GluN2A dependent neurotransmission affects functional links between neurons in the network. This detailed analysis of network parameters is critical in order to understand how network function is affected by a variety of interventions.

## 3. Discussion

In this report, we demonstrate the utility of a laser-scanning photostimulation (LSPS) technique for localized release of caged glutamate combined with Ca^2+^ fluorescence imaging in determining functional connections between neurons in culture, network properties arising from those connections, and the effects that interventions have on those properties. First introduced in 1993 (Callaway & Katz, 1993), LSPS of caged glutamate has been widely used to determine functional connections between neurons in brain slice preparations (Shepherd et al., 2005; Sturm et al., 2014) and *in vivo* (Noguchi et al., 2019). Typically, this technique has been combined with whole-cell voltage clamp to map the position, synaptic strength, and number of connections to a single post-synaptic neuron as a tool to understand microcircuit structure and function. The utility of this technique has been further expanded with the use of additional caged neurotransmitters (Rial Verde et al., 2008; Cabrera et al., 2018) and more recently of caged compounds in which photolysis depends on different light wavelengths (Amatrudo et al., 2015; Cabrera et al., 2017), thereby allowing specific stimulation of multiple species of caged compounds. Thus, LSPS is a powerful tool for describing the properties of microcircuit connections between neurons.

### 3.1. LSPS and Ca^2+^ imaging

Much less often, LSPS has been combined with Ca^2+^ imaging and when done so, Ca^2+^ imaging has typically been used to follow localized, subcellular increases in cytosolic Ca^2+^, typically within a single dendritic spine or group of adjacent spines (Schiller et al., 1998; Wei et al., 2001; Noguchi et al., 2011; Korkotian et al., 2014). Thus, combining LSPS with Ca^2+^ imaging allows spatially evoked increases of cytosolic Ca^2+^ to be observed in neurons. This capability was utilized in the present study to monitor the response of the neurons targeted for LSPS.

LSPS also allows mapping of functional connections in a neuronal network by monitoring the activity of neurons surrounding the neuron targeted for photolytic release of glutamate. One such approach is to use LSPS with neurons cultured on microelectrode arrays to study functional connections between the targeted presynaptic neuron and post-synaptic responding neurons within the array (Ghezzi et al., 2008). The advantage of this approach is that action potential activity in responding neurons can be directly measured. However, while it is possible to identify single neuronal activity, it is not possible to resolve the location of the responding neurons beyond the limitation imposed by the spacing of the electrodes within the array. As a result, it is difficult to produce a detailed spatial map of the microcircuitry of the neuronal network. In addition, at present, this approach is difficult to translate to an *in vivo* setting.

An alternative approach to mapping functional connections within a neuronal network is to combine LSPS with imaging techniques that monitor neuronal activity. First published by Nguyen et al. (2017) with highly enriched cultures of rat cortical neurons and adapted in the present study with mixed cultures containing glia and neurons, the activity of large numbers of neurons within a network can be monitored by observing increases in cytosolic Ca^2+^ that occur as a result of action potential bursting (Murphy et al., 1992; Robinson et al., 1993; Opitz et al., 2002). This approach has the advantage of probing the spatial organization of functional connections between neurons, independent of spontaneous synchronized Ca^2+^ activity, thereby allowing for a detailed description of the microcircuitry within a network. Additionally, this approach should be adaptable for use with voltage indicators and *in vivo* experimentation.

Though Nguyen et al. (2017) demonstrated the general utility of mapping functional connections with LSPS and imaging, the properties of the mapped connections were not elucidated. The present work shows that blockade of AMPA receptors does not change the number of neurons responsive to LSPS, the number of LSPS-induced links nor the stability of those links. Conversely, block of NMDA receptors with APV resulted in a decrease the number of neurons responsive to LSPS, the number of links and link stability. Qualitatively similar results were obtained when extracellular Mg^2+^ was increased. These data suggest connections revealed by this technique are due to NMDA but not AMPA receptor activation.

This finding might reflect the relative distribution of post-synaptic glutamate receptors. Although found in both soma and dendrites (Atkin et al., 2015), NMDA receptors appear to be more highly localized to the soma, whereas AMPA receptors tend to be located in the dendrites (Dodt et al., 1998; Smith et al., 2003). Another factor might be attenuation of dendritic signaling to the soma (Larkum et al., 1998; Jaffe and Carnevale, 1999), i.e. even if AMPA receptors in the dendrites of responding neurons are activated by synaptic transmission from a targeted neuron, the degree of activation at the soma would not be sufficient to induce bursts of action potentials needed to elicit measurable Ca^2+^ responses. Finally, glutamate-induced increases of intracellular Ca^2+^ in the soma of cultured neocortical neurons are largely dependent on Ca^2+^ influx via activation of NMDA receptors but unaffected by AMPA receptor block with CNQX (Wang et al., 2002). Irrespective of the mechanism, mapping neuronal connections by measuring Ca^2+^ responses induced by LSPS directly interrogates the contribution of NMDA receptors to connectivity between neurons in the network.

### 3.2. LSPS and Ca^2+^ imaging can be utilized to study perturbations in network properties

Another goal of the present study was to determine whether LSPS coupled with imaging could provide detailed information on how acute perturbations affect NMDA receptor-dependent network connectivity. Towards this end, the stability of network properties was investigated first. The data demonstrated that the number of targeted neurons responding to LSPS was relatively stable over the 30-40 min experimental period. While the total number of links, and thus the average number of links per neuron (*k*), showed a small but significant increase, there was little change in the average clustering coefficient (*C*). Individual links between targeted and responding neurons were likewise highly stable with over 80% being maintained during the repeated LSPS mapping segments. Furthermore, glutamate responsiveness, defined as the amplitude of the Ca^2+^ response of the neuron targeted for LSPS, an indirect measure of excitability, slightly, but significantly increased between mapping sessions. The weight of stable links, an indirect measure of the processes that include synaptic transmission, also slightly increased. Similar results were observed when the experimental period was extended to 60-70 minutes (not shown) and three mapping sessions were included in the experimental protocol. Thus, without intervention, baseline network properties were stable or slightly increased over time.

Of all the perturbations tested, the most dramatic effects on LSPS link properties were produced by addition of APV. NMDA receptor inhibition with this drug produced a decrease in the number of neurons responsive to LSPS and the number of links, as noted above. In addition, the amplitude of the Ca^2+^ responses in targeted neurons (glutamate responsiveness) decreased, as did the amplitude of the Ca^2+^ response in responding neurons (weight of stable links). Interestingly, the change in glutamate responsiveness of targeted neurons was only weakly correlated with the change in the weight of stable links. This analysis suggests that the effect of APV on the Ca^2+^ responses of responding neurons was not predominantly due to changes in the excitability of the neuron targeted by LSPS. Since the responding neuron is not directly stimulated, these results suggest that the predominant effect of APV on stable links is due to effects arising from synaptic transmission.

To modestly perturb network properties, we used inhibition and positive modulation of GluN2A receptors, a NMDA receptor subtype previously been shown to play a significant role in excitatory synaptic transmission (Sanz-Clemente et al., 2013) and in functional connections between cortical neurons in culture (Patel et al., 2014). The GluN2A receptor blocker, NVP-AAM077, decreased the number of links, consistent with previous reports (Patel et al., 2014), without a significant effect on the number of neurons responsive to LSPS, thereby decreasing *k* and *C*. In addition, the glutamate responsiveness of targeted neurons was significantly decreased, and the weight of stable links between targeted and connected neurons was reduced, consistent with a reduction of neuron excitability, but also hinting at a reduction in synaptic transmission. In contrast to these results, the GluN2A positive modulator, GNE-0723, increased the number of links, *k*, *C*, glutamate responsiveness and weights of stable links. These effects would be anticipated for an intervention that increases the activity of a NMDA receptor subtype which is localized both in the soma and dendritic spines (Atkin et al., 2015). Altogether, these data present a consistent description for how modulation of NMDA receptor function affects network properties.

### 3.3 Summary

The utility of combining LSPS and Ca^2+^ imaging in defining properties of NMDA receptor-dependent connectivity in networks of cultured cortical neurons has been demonstrated. This approach allows investigation of properties of individual connections between neurons as well as mapping network connectivity by targeting large numbers of neurons with LSPS. Connectivity properties are stable enough over sufficient time periods such that the effect of interventions having acute effects on either neuronal excitability and/or synaptic transmission can be studied at the network level. Although the present study is limited to acute effects on excitatory neuronal networks in culture, this combination of techniques should be applicable to neuronal networks that include GABAergic neurotransmission, longer time periods and monitoring of network properties *in vivo*.

## 4. EXPERIMENTAL PROCEDURE

### 4.1. Mixed Cortical Cell Culture

Cortical cells were isolated from E17/18 Sprague-Dawley rat embryos, and digested with 0.25% trypsin (Invitrogen, Carlsbad, CA) and DNase I (Sigma, St. Louis, MO) for 30 minutes at 37 ºC. Digestion was deactivated with fetal bovine serum (Invitrogen, Gaithersburg, MD) in Ca^2+^-free and Mg^2+^-free Hank’s Balanced Salt Solution (Gibco, Gaithersburg, MD). Cells were filtered with 70 µm and 40 µm filters sequentially, counted, and diluted in supplemented neurobasal (NB) medium consisting of Neurobasal Medium (Invitrogen, Gaithersburg, MD), 2% B27, 1% penicillin–streptomycin, and 0.4 mM L-glutamine, as previously published (Magou et al., 2015). All procedures were approved by the Newark Institutional Animal Care and Use Committee at Rutgers University.

Cultures were seeded at an average density of 110,000 cells per cm^2^ on a 0.125 mm (0.005 in.) thick elastic silicone substrate (Specialty Manufacturing Inc., Saginaw, MI) in custom 16-mm diameter wells that were assembled, autoclaved and dried in sterile conditions, as described previously (Pfister et al., 2003). Prior to plating, the silicone substrate was incubated with 40 µg/ml poly-L-lysine hydrobromide (Sigma, St. Louis, MO) dissolved in sterile distilled water overnight, then rinsed with PBS three times, and dried overnight at room temperature. Cultures were maintained in supplemented NB medium at 37 ºC with 5% CO2 for 18-22 DIV when cultures display periodic bursting electrical activity (Kamioka et al., 1996), a sign of a highly developed neuronal network (van Pelt et al., 2004).

### 4.2.Fluorescence Imaging of Spontaneous Network Activity

Prior to experiments, cells were loaded with the calcium indicator Fluo-4 by incubating in supplemented NB medium that contained 2 µM of Fluo-4 AM (Invitrogen, Carlsbad, CA), and Pluronic F-127 (0.04%) for 30 minutes, then washed with artificial cerebral spinal fluid (ACSF) for an additional 10 minutes to allow for de-esterification of Fluo-4. The ACSF used as the bath solution for all experiments, was comprised of (in mM): 140 NaCl, 24 D-glucose, 10 HEPES, 5 KCl, 1.8 CaCl2 and 1 MgCl2; pH 7.2 with NaOH. Bicuculline methiodide (40 µM) was added to ACSF solutions during the washes and experiments unless otherwise noted. All experiments were performed at room temperature, approximately 25°C.

Fluo-4 loaded cultures were placed under an upright microscope at 4X magnification with a field-of-view (FOV) area of 1350 µm by 1350 µm, containing 600-1000 neurons, and illuminated by 494-nm light from a high-power LED. Emission fluorescence at 516 nm (50 nm bandwidth) wavelength was captured at 1 Hz with an EMCCD (ImagEM X2, Hamamatsu, Bridgewater, NJ). A custom-written LabVIEW (National Instruments, Austin, TX) program interfacing with ImageJ (NIH, Bethesda, MD) allowed for fluorescent image stacks of spontaneous Ca^2+^ activity to be automatically captured and saved.

### 4.3. Pharmacological Agents

Where indicated, drugs were added as a 10-20 µl addition of stock solution into the ACSF bath and 10 minutes were allotted for the drug to mix throughout the solution. Final drug concentrations were 40 µM bicuculline methiodide, 2 µM 6-Cyano-7-nitroquinoxaline-2,3-dione (CNQX), 25 µM 2-Amino-5-phosphono-pentanoic acid (APV), 50 nM NVP-AAM077 or 1 µM GNE-0723 (kindly provided by Genentech, Inc, San Francisco, CA). Unless, otherwise indicated, drugs were purchased from Sigma-Aldrich, Inc. (St. Louis, MO).

### 4.4. Laser Scanning Photostimulation (LSPS) to Map Network Connections

In these experiments, MNI-caged glutamate (Tocris Bioscience, UK), added to a final concentration of 200 µM, was photolyzed in a region surrounding the soma of a single neuron using light from a 375-nm diode laser (Power Technology, Alexander, AR) set at 2 mW power. The light was focused through the 4x objective to a 10-µm spot and current-modulated to produce a 2-ms pulse (Nguyen et al., 2017). Stimulated activities of responding cells were recorded as increases in Fluo-4 fluorescence in images captured at 10 Hz with 50-ms exposures over a period of 2.5 seconds after the laser pulse. Each LSPS session targeted 75 neurons and lasted for approximately 10 minutes. A custom-written LabVIEW program enabled the LSPS protocol to be automated with little user intervention. Detection and analysis of Ca^2+^ signals were performed using custom-written MATLAB (MathWorks, Natick, MA) routines.

### 4.5. Statistical Analysis

Tests for normality and equal variance were performed for all data groups. Those data satisfying these tests are displayed as mean ± S.E.M., as indicated. Otherwise, data are displayed as median and magnitude of the interquartile range (IQR), i.e. 25-75% quartile range, in text, tables and graphs. For nonparametric analyses, comparisons between experimental groups were made with Mann-Whitney U or Wilcoxon Signed Ranks tests. A statistically significant change was defined by p < 0.05. Statistical tests were performed in R or Prism (GraphPad Software, San Diego, CA).

## Abbreviations

LSPS: laser scanning photostimulation
ENN: Excitatory-only neuronal network
*k*: average degree
*C*: average clustering coefficient

## ACKNOWLEDGEMENTS

The authors wish to thank Bogumila Swietek and Dr. Ying Li for isolation for cortical tissue and assistance with immunohistochemistry, Nirali Trivedi and George Mina for assistance with cell cultures, Mina Ghbrial for assistance with data analysis, and Olivia Ortelli for assistance with network analysis.

